# Synthetic Community Inoculation on Seeds Revealed its Transient Colonization Capacity but Legacy Effects on Plant Microbiota Assembly

**DOI:** 10.1101/2025.03.28.645722

**Authors:** Gontran Arnault, Louna Colaert-Sentenac, Coralie Marais, Alain Sarniguet, Matthieu Barret, Marie Simonin

**Author notes:** **Corresponding author**: Gontran Arnault.

## Abstract

Seed microbiota have the potential to influence the overall plant microbiota assembly. However, to date, studies have mostly focused on early plant development stages. This study aimed to investigate the influence of seed microbiota on plant microbiota assembly throughout an entire life cycle. To achieve this, bacterial synthetic communities (SynComs) were reconstructed and inoculated on common bean seeds to eliminate the natural variability of seed microbiota and provide four different primary inocula for comparison. We then examined bacterial and fungal communities at different developmental stages (seedling, vegetative, flowering, pod-filling, and senescent stages) and in different plant compartments (rhizosphere, root, leaf, seed) of the common bean. SynComs inoculated on seeds significantly contributed to the seedling microbiota, with higher colonization success in the leaves compared to roots and rhizospheres. Strain identity and SynCom composition influenced the strain colonization capacity across the habitats. Also, bacterial SynCom colonization induced composition modification in the seedling root and leaf microbiota. After the seedling stage, SynComs members were not detected in plant compartments but promoted persistent changes in microbial community composition until the next generation of seeds. In conclusion, SynCom inoculated on seeds have a transient colonization that can influence the overall plant microbiota assembly through priority effects.

## Introduction

Plants harbor specific microbiota in each compartment, which include among others the rhizosphere (zone of soil surrounding and influenced by the root), the root, the leaf and the seed microbiota (Xiong et al., 2021). These compartments are assembled from many environmental and maternal sources such as soil, air, insect, seed or clonal organ (Lundberg et al., 2012; Prado et al., 2020; Nelson 2018, Rezki et al., 2018; Vannier et al., 2018). Soil is often described as the main source of microorganisms and its origin highly influences the plant microbiota assembly (Xiong et al., 2021; Lundberg et al., 2012; Schlaeppi et Bulgarelli, 2015; Fierer 2017; P. Brown et al., 2020). However, the potential transmission through seeds has been less investigated as most of the plant microbiota assembly studies use surface sterilized seeds and/or do not describe this potentially important source compartment (Xiong et al., 2021; Lundberg et al., 2012, Nelson 2018). The seed microbiota is nevertheless the primary inoculum of plant microbiota and could thus influence the overall plant microbiota assembly through priority effects (Abdelfattah et al., 2022; Debray et al., 2021; Ridout et al., 2019; Arnault et al., 2024). Thus, more effort is needed to better understand how the seed microbiota might influence the overall plant microbiota assembly (Nelson 2018).

Seed is a key organ of the spermatophyte life cycle. During the seed-to-seed cycle, seeds can be colonized by many microorganisms such as bacteria or fungi (Nelson 2018). These microorganisms can be vertically transmitted to seed through the vascular pathway, or horizontally acquired from the environment during seed development (floral pathway), seed dispersal and contact with the soil (Nelson 2018, Abdelfattah et al., 2022). The seed microbiota is influenced by production site, plant species and plant genotype (Rochefort et al., 2019; Simonin et al., 2022; Malacrino et al., 2022). A specificity of seeds is that microbial biomass (number of cells) and richness (number of species) are reduced compared to other plant compartments (War et al., 2023; Guo et al., 2021). Additionally, seed microbiota biomass and composition are highly variable from one seed to another and most seeds are dominated by a single taxon (Ridout et al., 2019; Kim et al., 2023; Chesneau et al., 2022). Consequently, the few studies which have analyzed the importance of seeds as a potential source for plant microbiota assembly present contrasting results depending on plant species and experimental designs (Rochefort et al, 2021; Walsh et al, 2021, Johnston Monje et al., 2016; Chesneau et al., 2022). Some reported a dominance of soil borne taxa over seed microbiota (Rochefort et al., 2021; Walsh et al., 2021) while some others reported a dominance of seed borne taxa on plant microbiota assembly (Johnston Monje et al., 2016; Johnston Monje et al., 2021; Moroenyane et al., 2021). The main limitation of these studies is the use of seed batches to describe the initial seed source, while individual seeds are known to have very distinct microbiota. Moreover, most studies focus solely on early stages of plant development (Simonin et al. 2023, Rochefort et al., 2021, Chesneau et al., 2022, Arnault et al., 2024, Johnston-Monje 2021, Walsh et al., 2021), although a few have described seed-to-seed transmission of microorganisms, but still in seed batches (Rezki et al., 2018; Kim et al., 2022; Bergna et al., 2018). The importance of single seed microbiota as a potential inoculum for the plant microbiota assembly still faces technical limitations and need to be further studied.

One possible solution to control the natural variation of the seed microbiota is the use of Synthetic Community (SynCom) to establish causality between seed microbiota composition and plant microbiota assembly (Vorholt et al., 2017, Mehlferber et al., 2024, Arnault et al., 2024). Some studies reported that the use of SynComs on seed could modify both seed and seedling microbiota assembly (Simonin et al., 2023, Arnault et al., 2024). Our studies showed that SynCom on seeds can manipulate 97% of the relative abundance of the seed microbiota and 80% of the seedling microbiota, and could influence the bacterial microbiota assembly of both seedling and rhizosphere (Arnault et al., 2024). However, the distinct seed routes from seed to seedling are not well elucidated and were not studied in our previous studies (Abdelfattah et al., 2022, Simonin et al., 2023, Arnault et al., 2024). Also, to the best of our knowledge, the inoculation of SynCom on seed to study the influence of seed microbiota composition on plant microbiota assembly across the entire plant life cycle has never been reported before.

In this study, we manipulated the seed microbiota using SynComs to study its colonization success and influence on leaf, root, rhizosphere and seed microbiota assemblies. To do so, 22 bacterial isolates from common bean seeds were used and 3 different SynComs of contrasted composition were inoculated. We monitored each individual bacterial strains and the assembly of the bacterial and fungal communities using amplicon sequencing at 5 phenological stages. This study aims to address two unanswered questions: (i) What is the colonization success of the seed microbiota members during plant development? (ii) What influence do they have on plant microbiota assembly across plant stages and compartments?

## Material and methods

### SynCom design and inoculation

This research aimed to investigate plant microbiota assembly using different compositions of seed microbiota as starting points. Each SynCom was composed of 8 strains using a pool of 22 strains isolated from common bean seeds (Arnault et al., 2024).

The experiment was performed using a commercial seed lot of Flavert genotype from Vilmorin-Mikado® (France). Seeds were rinsed three times using sterilized water before inoculation. SynComs were inoculated at 10^7^ CFU/mL using water and by scratching a 48 h culture of TSA 10% for each strain, which was adjusted to an OD (600 nm) that matches a population size of 10^7^ CFU/mL. SynComs were prepared by adding equivolume of each strain. Bacterial population and SynCom biomass (measured in colony forming units - CFU) of the inocula were verified by dilution and plating on TSA 10% and incubation at 18°C for 4 days. SynComs inoculations were performed by placing the seeds in a sterile container and adding 2 mL of inoculum (sterile water for control) per gram of seed for 30 min under constant agitation (70 rpm) at 18°C. Excess inocula were then removed using a sterile strainer, and seeds were dried for 30 min under a laminar flow. Seed bacterial community biomass was measured after inoculation on 8 seeds per condition. To do so, each seed was soaked in 2 mL of sterile water at 4°C under constant agitation (220 rpm) overnight. The resulting suspensions were then plated on TSA 10% and microbiota profiling was performed on the same 8 biological replicates.

### Plant growth and DNA extraction

Inoculated seeds (n=50 per SynCom) were sown in a non-sterile potting soil (Traysubstrat Klasmann-Deilmann France) in a greenhouse (16 h day at 23°C, 8 h night at 20°C, 70% humidity; Figure 1). Plants were harvest at 5 different phenological stages hereafter called: (i) seedling [5 Days After Sowing (DAS), BBCH stage 12, two full leaves unfolded], (ii) vegetative stage (20 DAS, BBCH stage 14, 4 trifoliate), (iii) flowering stage (34 DAS, BBCH stage 55, first flower buds enlarged), (iv) podfilling stage (54 DAS, BBCH stage 79) and (v) senescence stage (84 DAS, BBCH stage 99). For seedling, vegetative, flowering and podfilling stages, the following plant compartments were sampled: rhizosphere, root, shoot, leaf. For the flowering stage, flower buds were also sampled. Most of the shoot and flower bud samples were lost during library preparation due to lack of microbial biomass (no amplification) and were thus removed from the analysis. For podfilling and senescence stages, seeds were also harvested by taking 1 seed at random from 25 pods also picked randomly (pool of 25 seeds analyzed). The adhering soil to plant roots was considered as rhizosphere and conserved at - 80°C before DNA extraction. For each development stage, primary roots were sampled and rinsed twice using sterilized water, cleaned using sterile brushes and crushed with a roller. Then, 5mL of sterilized water was added and the roots were ground for 30 s using a stomacher. For the seedling stage, the two first leaves were harvested and crushed with a roller. Then 2 mL of sterilized water was added and the leaves were ground for 30 s using a stomacher. For the vegetative stage, the first 3 trifoliates were harvested and crushed with a roller. For the flowering and podfilling stages, two leaflets of each trifoliates were harvested and crushed with a roller. Then 40 mL of sterilized water was added and the leaves were ground for 30 s using a stomacher. The suspension obtained was centrifugate 10mn at 4000g and the pellet was resuspended in 1mL of sterilized water and stored at -80°C before DNA extraction.

**Figure 1:**
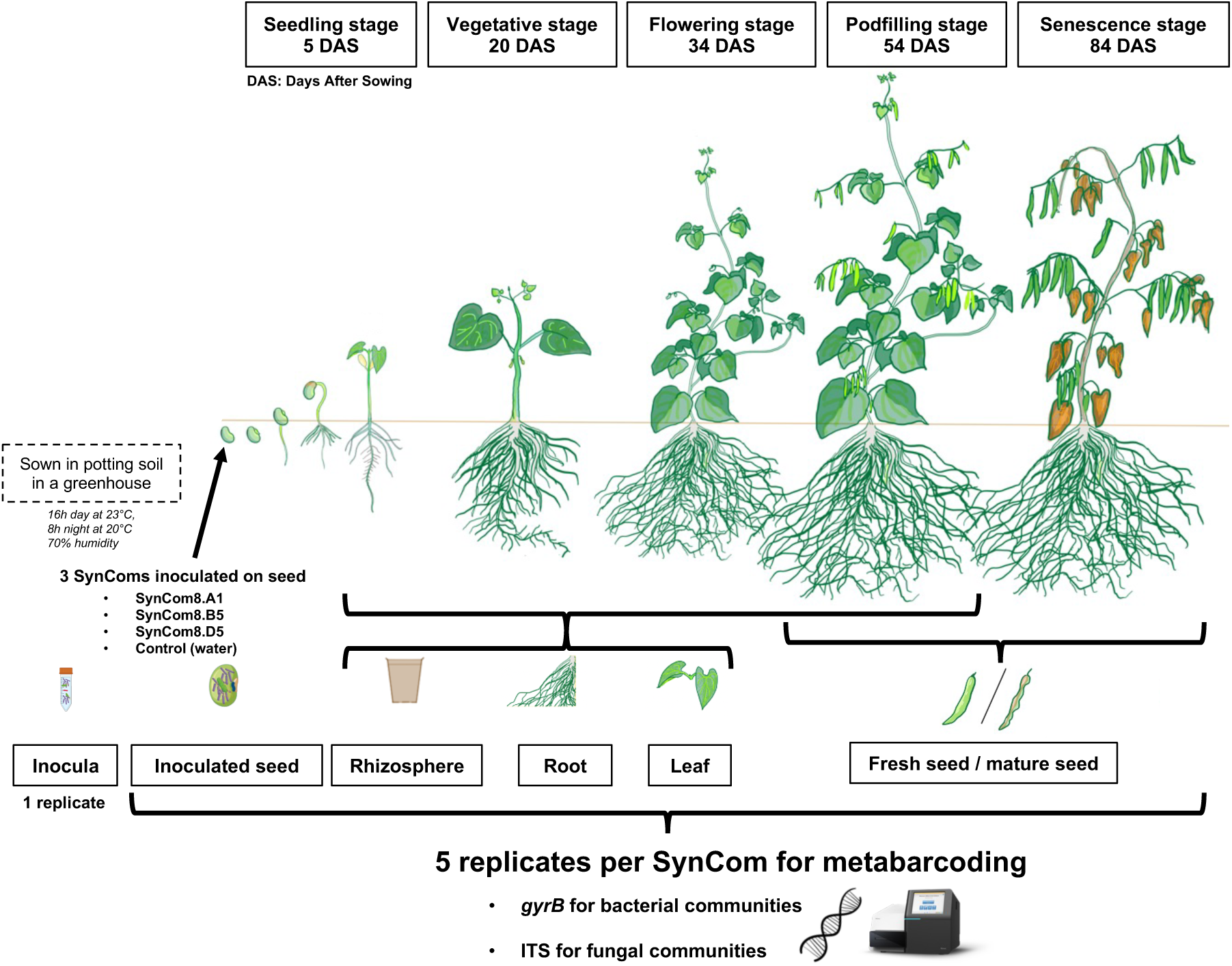
experimental design of common bean sampling after SynCom inoculation on seed. Three SynComs were inoculated on seeds (water for control). Seeds were sown in non-sterile potting soil in a greenhouse with these parameters: 16h day (23°C), 8h night (20°C), 70% humidity. Plants were sampled at 5 different plant development stages: (i) seedling [5 Days After Sowing (DAS), BBCH stage 12, two full leaves unfolded], (ii) vegetative stage (20 DAS, BBCH stage 14, 4 trifoliate), (iii) flowering stage (34 DAS, BBCH stage 55, first flower buds enlarged), (iv) podfilling stage (54 DAS, BBCH stage 79) and (v) senescence stage (84 DAS, BBCH stage 99). Rhizosphere, leaf, shoot, root, flower bud and seeds were sampled for each appropriate development stage. Five plant replicates were analyzed by metabarcoding for bacterial (gyrB) and fungal (ITS1) community analyses.

### Metabarcoding on *gyrB* and ITS genes and taxonomic classification

The following metabarcoding approach was performed using 200 μL of each suspension described before. For inocula, 200 μL of each fresh inoculum was instantly stored at -80°C before DNA extraction. DNA was extracted using the NucleoSpin® 96 Food kit (Macherey-Nagel, Düren, Germany) following the manufacturer’s instructions. For rhizosphere characterization, ∼200 mg per condition was extracted using DNA PowerSoil kit from Qiagen following the manufacturer’s instructions. Initial potting soil community composition (day of sowing) was also characterized using the same procedure as the rhizosphere and using 4 replicates.

The first PCR was performed with the primers gyrB_aF64/gyrB_aR553 (Barret et al. 2015), which target a portion of bacterial *gyrB* gene, and ITS1F/ITS2 primers (Buée et al., 2009), which target fungal ITS1 region. PCR reactions were performed with a high-fidelity Taq DNA polymerase (AccuPrimeTM Taq DNA polymerase Polymerase System, Invitrogen, Carlsbad, California, USA) using 5 µL of 10X Buffer, 1 µL of forward and reverse primers (100 µM for *gyrB* and 10 µM for ITS1), 0.2 µL of Taq and 5 µl of DNA. For gyrB_aF64/gyrB_aR553 primers, PCR cycling conditions were done with an initial denaturation step at 94°C for 3 min, followed by 35 cycles of amplification at 94°C (30 s), 55°C (45 s) and 68°C (90 s), and a final elongation at 68°C for 10 min. For ITS1F/ITS2 primers, the cycling conditions were an initial denaturation at 94°C for 3 min, followed by 35 cycles of amplification at 94°C (30 s), 50°C (45 s), and 68°C (90 s), and a final elongation step at 68°C for 10 min. Amplicons were purified with magnetic beads (Sera-MagTM, Merck, Kenilworth, New Jersey). The second PCR was conducted to incorporate Illumina adapters and barcodes. The PCR cycling conditions were the same for the two markers: denaturation at 94°C (2 min), 12 cycles at 94°C (1 min), 55°C (1 min) and 68°C (1 min), and a final elongation at 68°C for 10 min. Amplicons were purified with magnetic beads and pooled. Concentration of the pool was measured with quantitative PCR (KAPA Library Quantification Kit, Roche, Basel, Switzerland). Amplicon libraries were mixed with 15% PhiX and sequenced with three MiSeq reagent kits v2 500 cycles (Illumina, San Diego, California, USA). A blank extraction kit control, a PCR-negative control and PCR-positive control (*Lactococcus piscium* DSM6634, a fish pathogen that is not plant-associated) were included in each PCR plate.

The bioinformatic processing of the amplicons was performed in R. Primer sequences were removed with cutadapt 2.7 (Martin, 2011) and trimmed fastq files were processed with DADA2 version 1.22.0 (Callahan et al., 2016). Chimeric sequences were identified and removed with the removeBimeraDenovo function of DADA2. For fungi, only forward reads were used due to the bad quality of the reverse reads. Amplicon Sequence Variant (ASV) taxonomic affiliations were performed with a naive Bayesian classifier (Wang et al. 2007) with our in-house *gyrB* database (train_set_gyrB_v5.fa.gz) for bacteria and using UNITE V8 database for fungi. The post-clustering algorithm LULU was used to identify SNPs and merge them using a clustering level of 98% for bacteria and 99% for fungi. Then, the identification of sequence contaminants was assessed with decontam v1.20.0 using the prevalence method (threshold = 0.5 for bacteria, 0.2 for fungi) and some contaminants were also removed by manually checking control sample compositions. Unassigned sequences at the phylum level and *parE* sequences (a *gyrB* paralog) were filtered for bacteria and sequences affiliated to Viridiplantae and Rhizaria were removed for fungi. To track our SynCom strains, only ASVs with 100% of identity were considered to be our strains.

### Statistical analyses and microbiota analysis

Microbial community analyses were conducted using the Phyloseq package v1.44.0 (Mc Murdie and Holmes 2013) and figures were generated using ggplot2 v3.4.3. A coverage-based rarefaction was made using iNEXT package v2.6.4 for both *gyrB* and ITS1 phyloseq objects (Hsieh et al. 2016). Wilcoxon tests were used to compare richness between SynCom inoculated microbiota and control as well as cumulative relative abundance of SynCom ASVs. Beta diversity analyses were made using BrayCurtis distance, PCoA ordination and permutational multivariate analysis of variance [adonis2 function of vegan v2.6.4 (Oksanen et al. 2019), 999 permutations]. Differential abundances testing of communities was evaluated employing a linear model through the CornCob package v0.4.1 using taxatree_models function (after a log2 transformation) and taxatree_plots function for visualization. Linear regression between seedling root and leaf strain relative abundances was obtained using lm function from stats package.

## Results

### Transient Colonization of the Three Seed Synthetic Communities

To study the colonization success of SynComs, we tracked our strains across plant development stages and compartments (Figures 2 & 3). The cumulative relative abundance of SynCom ASVs were compared with the control to assess SynCom colonization capacities. At the seedling stage, SynComs significantly colonized all compartments compared to control seedlings (Figure 2-A, Wilcoxon-test; p-values < 0.05). SynCom ASVs were more abundant in the leaf microbiota (56.1% to 94.7%) compared to the root (6 to 32.4%) and rhizosphere (0.3% to 14.8%). At the vegetative, flowering and podfilling stages, the cumulative relative abundance of SynComs ranged from 0 to 2.8% across the different compartments and were not significantly different from the control plants (Figure 2-A, Wilcoxon-test; p-values < 0.05).

**Figure 2:**
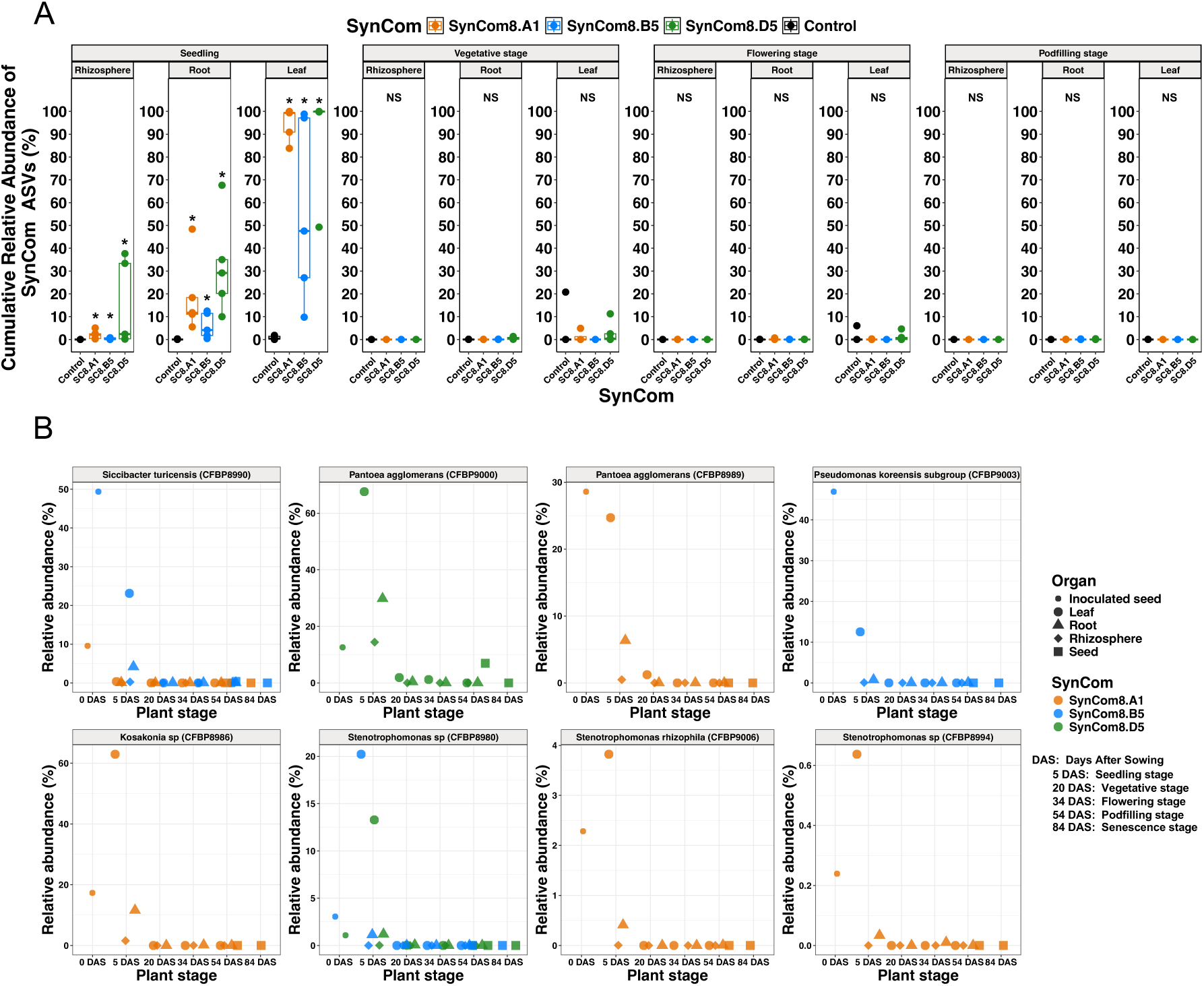
Colonization by the SynComs (A) and strains (B) of the different plant compartments across plant development stages. (A) Cumulative relative abundance of SynCom ASVs across plant development stages and compartments. Wilcoxon tests were used to compare each SynCom with the control (*: p-values < 0.05; NS: Not significant). (B) Relative abundance of 8 strains across development stages and plant compartment.

**Figure 3:**
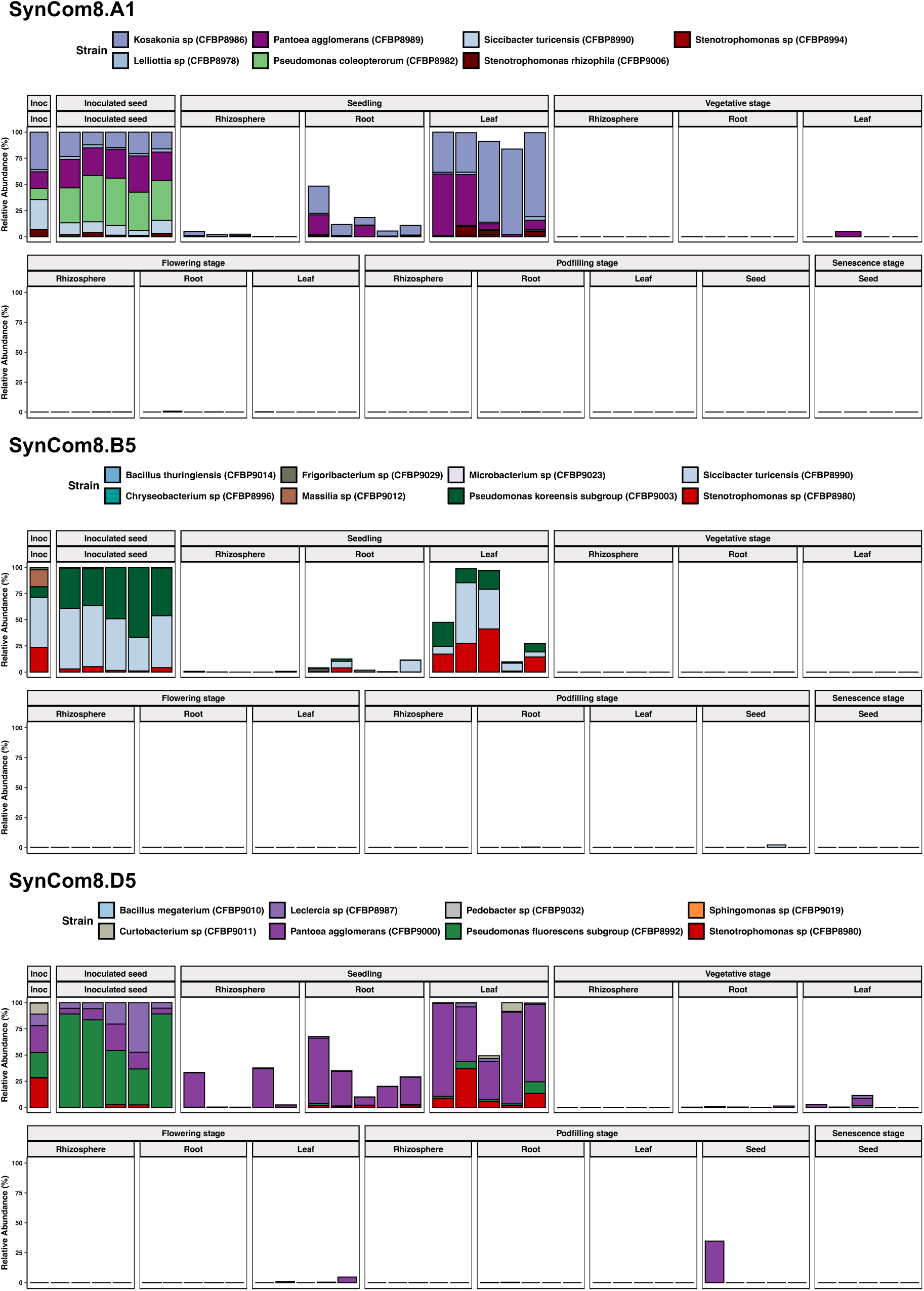
SynCom taxonomic profiles across plant stages and compartments. Taxonomic profiles of each SynCom across inocula, inoculated seed, seedling, vegetative, flowering, podfilling and senescence stages.

Individual strain transmission capacities were also analyzed across compartments and plant stages (FigS1, Figure 2-B & Figure 3). Strains presented highly contrasted transmission capacities (FigS1). For instance, *Pedobacter sp* (CFBP9032), *Chryseobacterium sp* (CFBP8996), *Bacillus thuringiensis* (CFBP9014) and *B. megaterium* (CFBP9010) were never transmitted even to seedlings while 8 strains were always transmitted to seedlings in both leaf and root (FigS1). A subset of the high seedling colonizers visible in the taxonomic profiles is presented on Figure 2-B. The relative abundance of these strains across plant development stages highlighted their slow disappearance from inoculated seed to mature seed of the next generation (Figure 2-B). Also, all strains were better leaf colonizers than root and rhizosphere colonizers (Figure 2-B). From seed to seedling, 5 strains (CFBP8980, CFBP8986, CFBP8994, CFBP9000 & CFBP9006) increased their relative abundance while 3 strains (CFBP8989, CFBP8990 & CFBP9003) decreased their relative abundance. Thus, strains show fitness differences between the seed and the seedling niches. *Siccibacter turicensis* (CFBP8990) had a transmission rate of 80% in rhizospheres of SynCom8.B5 while it was never transmitted to rhizospheres of SynCom.A1. For seedling root and leaf, this strain was always transmitted for SynCom8.B5 while it had a transmission rate of 60% in SynCom.A1. These results highlight that SynCom composition can influence individual strain transmission from seed to plant. For the well-transmitted strains (relative abundance > 0.05%, n = 14), their relative abundance on leaf was correlated with their relative abundance on root (FigS2; R^2^ = 80%; p-value < 0.001). Thus, even if SynCom strains represent a higher relative abundance on leaf than root, the colonization success between these two niches is correlated.

In conclusion, SynComs ASVs represented a high proportion of the seedling leaf microbiota. They also colonized seedling roots and rhizospheres to a lesser extent. After the seedling stage, their transmission rates were very variable and their relative abundance very low indicating a transient colonization of these seed-borne strains. Strain identity and SynCom composition influenced the seed to plant colonization success.

### Impact of synthetic community inoculation on plant microbiota assembly

Next, we assessed the impact of this transient colonization of the three SynComs on the assembly of bacterial and fungal communities across plant development stages.

For the bacterial communities, rhizosphere and root alpha diversity presented the same dynamic: seedlings had less richness (345 and 385 ASVs respectively) than the other development stages. Then, a high increase in richness occurred at vegetative stage with 612 ASVs in the rhizosphere and 704 ASVs in the root. Next, bacterial diversity slowly decreased until podfilling stages (496 ASVs in the rhizosphere and 589 ASVs in the root). Globally, root bacterial communities presented more observed richness than the rhizosphere. Leaf bacterial communities were less diverse than both rhizosphere and root communities and fluctuated according to plant stages (ranging from 66 ASVs at vegetative stage to 195 ASVs at flowering stage). For fungi, unlike bacteria, the rhizospheres were more diversified (from 66 to 78 ASVs) than the root microbiota (from 23 to 34 ASVs) and both presented stable alpha diversity across development stages. Fungal leaf microbiota were more variable with a mean of 22 ASVs at seedling stage, 67 ASVs at vegetative and 62 ASVs at flowering stages and had an increase to 130 ASVs at podfilling stage. Globally, the rhizosphere and root microbiota presented more bacterial diversity compared to fungal diversity. Leaf microbiota had more bacterial diversity at seedling stage but more fungal diversity at podfilling stage.

SynCom inoculation on seed did not impact either bacterial (Fig4-A) nor fungal (Fig4-B) alpha diversity across the different plant development stages and compartments compared to control seeds (Figure 4). Thus, the development stage and compartment drove alpha diversity, whereas the seed microbiota composition did not.

**Figure 4:**
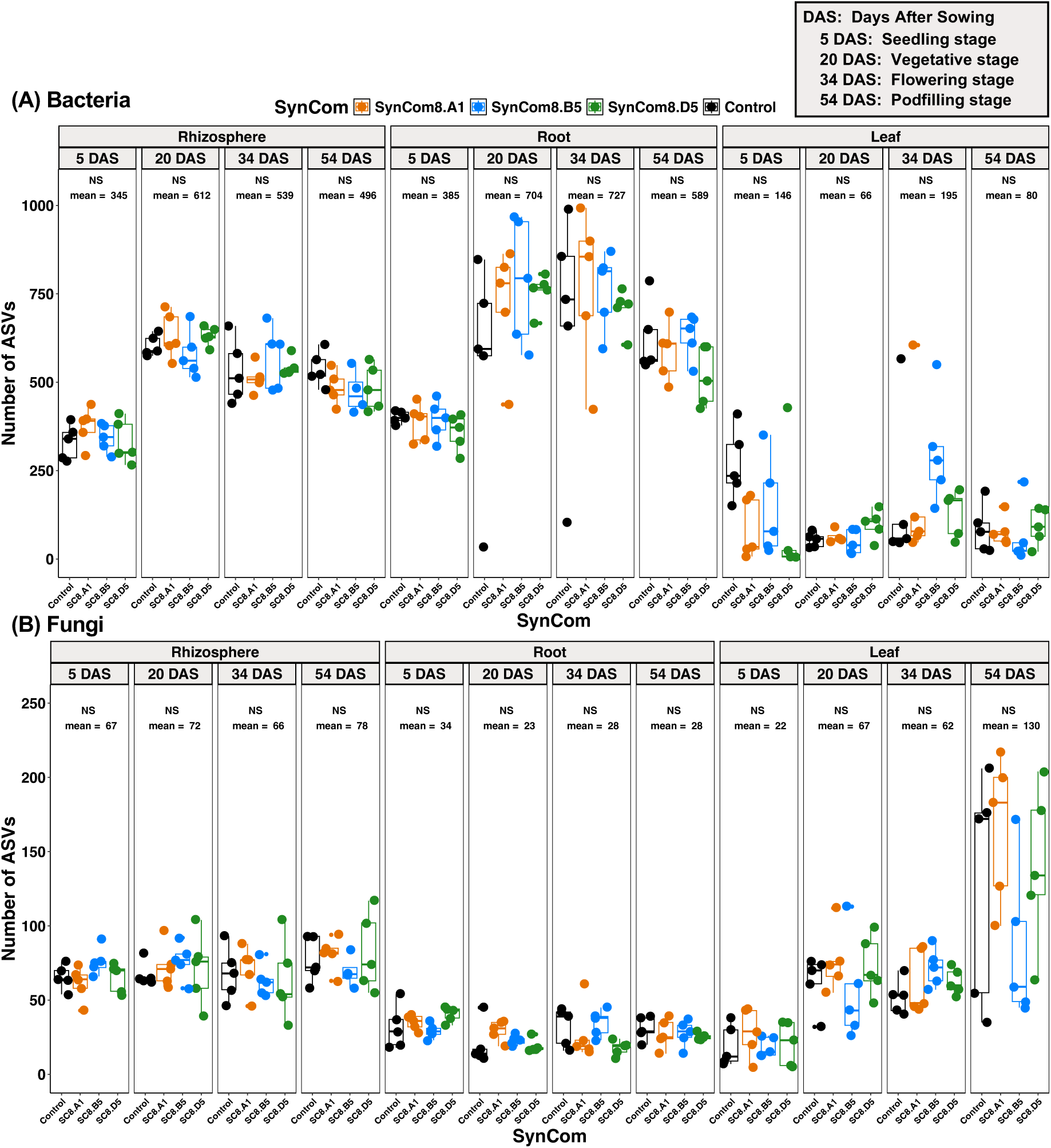
Alpha diversity across plant development stages and compartments for bacterial (A) and fungal (B) communities. Alpha diversity was measured as the number of bacterial (A) and fungal (B) ASVs across plant development stages and compartments. Development stages are indicated as Days After Sowing (DAS). 5 DAS: seedling stage (BBCH stage 12, two full leaves unfolded); 20 DAS: vegetative stage (BBCH stage 14, 4 trifoliate); 34 DAS: flowering stage (BBCH stage 55, first flower buds enlarged); 54 DAS: podfilling stage (BBCH stage 79). Wilcoxon-tests were used to compare the control with each SynCom for each combination of compartment and development stage (NS: non significant). Mean number of ASVs across conditions for each combination of compartment and development stage are indicated on the plot.

Bacterial community structures were then analyzed using PCoA ordination based on Bray-Curtis distances and PERMANOVA analyses. SynCom inoculation drove the initial seed microbiota composition (Figure 6, PERMANOVA; R^2^ = 69.8%, p-value = 0.001). All seed conditions (3 SynComs + control) had significantly different compositions, which initiated the plant microbiota assembly with 4 different seed microbiota compositions. Bacterial rhizosphere, root and leaf communities were primarily structured by plant stage (Figure 5-A, PERMANOVA; R^2^ = 31.3%, 23.2%, and 25.9% respectively, p-values < 0.001). Rhizospheres were dominated by *Burkholderiales*, *Micrococcales*, *Rhizobiales* and *Xanthomonadales*, root communities by *Burkholderiales*, *Rhizobiales* and *Myxococcales.* Leaf communities presented more variability in taxonomic composition, but with a dominance of *Burkholderiales* at podfilling stage. The effect of SynCom on bacterial microbiota assembly was only significant in leaf (Figure 5-A, PERMANOVA; R^2^ = 5.2%, p-value = 0.002). A significant interaction between plant development stage and seed inoculation of SynCom was observed for this habitat (PERMANOVA; R^2^ = 15.5%, p-value = 0.001). By analyzing dissimilarity in bacterial community composition within each plant development stage, SynCom inoculation was the main driver of seedling leaf community structure (Figure 6, R^2^ = 55.4%, p-value < 0.001). This effect is also found to a lesser extent for bacterial communities associated with seedling roots (Figure 6, PERMANOVA; Root: R^2^ = 30.6%, p-value < 0.001). The effect of SynComs inoculation on bacterial community structure was neither observed at the following development stages (i.e. vegetative, flowering and podfilling stages) nor to the next seed generation (i.e. fresh and mature seeds). For seedling leaf microbiota, all bacterial communities showed significantly different composition. SynCom inoculation explained from 28.1% (control *versus* SynCom8.B5 leaf communities) to 63.5% (SynCom8.A1 *versus* SynCom8.D5 leaf communities) of the leaf microbiota composition variance (Pairwise permanova, p-values < 0.05). For seedling root microbiota, all SynComs presented significantly different compositions between each other (Pairwise permanova, p-values > 0.05). SynCom inoculation on seed did not impact seedling rhizosphere microbiota composition (PERMANOVA; R^2^ = 19.7%, p-value = 0.143). For the vegetative, flowering and podfilling stages, initial seed SynComs had no influence on bacterial community composition (Figure 6, PERMANOVA; p-values > 0.05). However, many bacterial genera were enriched or depleted across plant stages and compartments according to initial SynCom composition (Figure 9-A). Seed microbiota of the next generation was investigated at the podfilling (fresh seeds, 54 DAS) and senescence stages (mature seeds, 84 DAS) (Figure 5-B & 6). Also, initial seed composition post SynCom inoculation did not influence the overall microbiota structure of the next generation seeds (Figure 6). Fresh seeds were mainly composed of *Burkholderiales* and *Enterobacterales.* Mature seed compositions were more variable at order level, with mostly *Burkholderiales, Enterobacterales, Sphingomonadales.* However, differential abundance testing revealed that mature seeds from control condition were enriched in *Burkholderiales*, specifically in the genus *Herbaspirillum*, mature seeds from SynCom8.A1 were enriched in *Enterobacterales* specifically in the genus *Kluyvera* and mature seeds from SynCom8.D5 were enriched in *Bradyrhizobium* (Figure 9-A and FigS3). Thus, the SynCom inoculation had an influence on the seed composition of the next generation.

**Figure 5:**
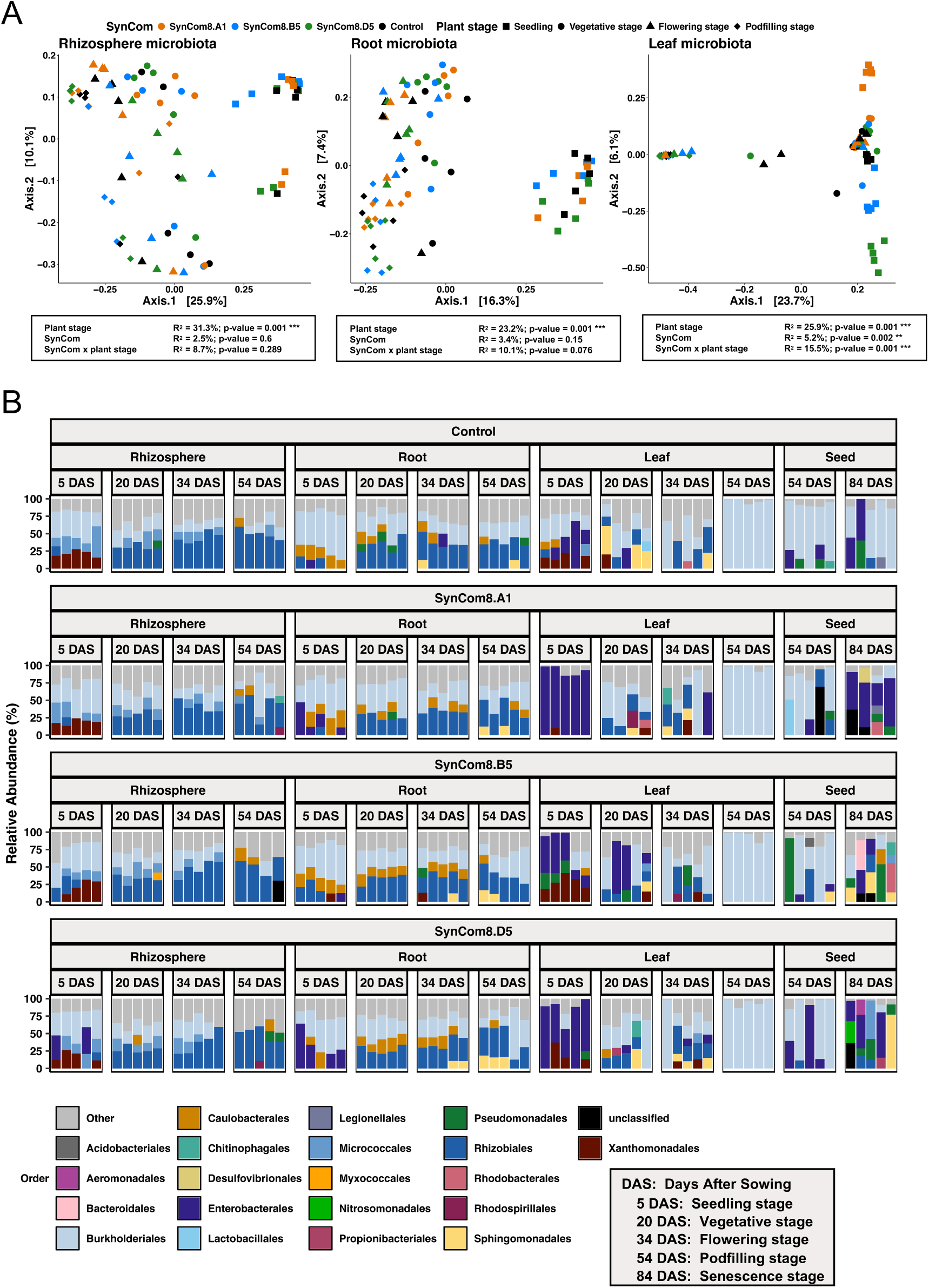
Overview on bacterial communities across plant stages and compartments. (A) Effect of SynCom and plant development stages on rhizosphere, root and leaf bacterial microbiota structure visualized through PCoA ordinations based on Bray–Curtis distances. PERMANOVA were used to test the effect of plant development stage, SynCom and their interaction for each plant compartment. Samples are shaped based on their plant stage and colored by their initial SynCom composition. (B) Taxonomic profiles at order level of the plant compartments across stages and SynComs. Taxa representing less than 10% of relative abundance were merged in the “Other” category.

**Figure 6:**
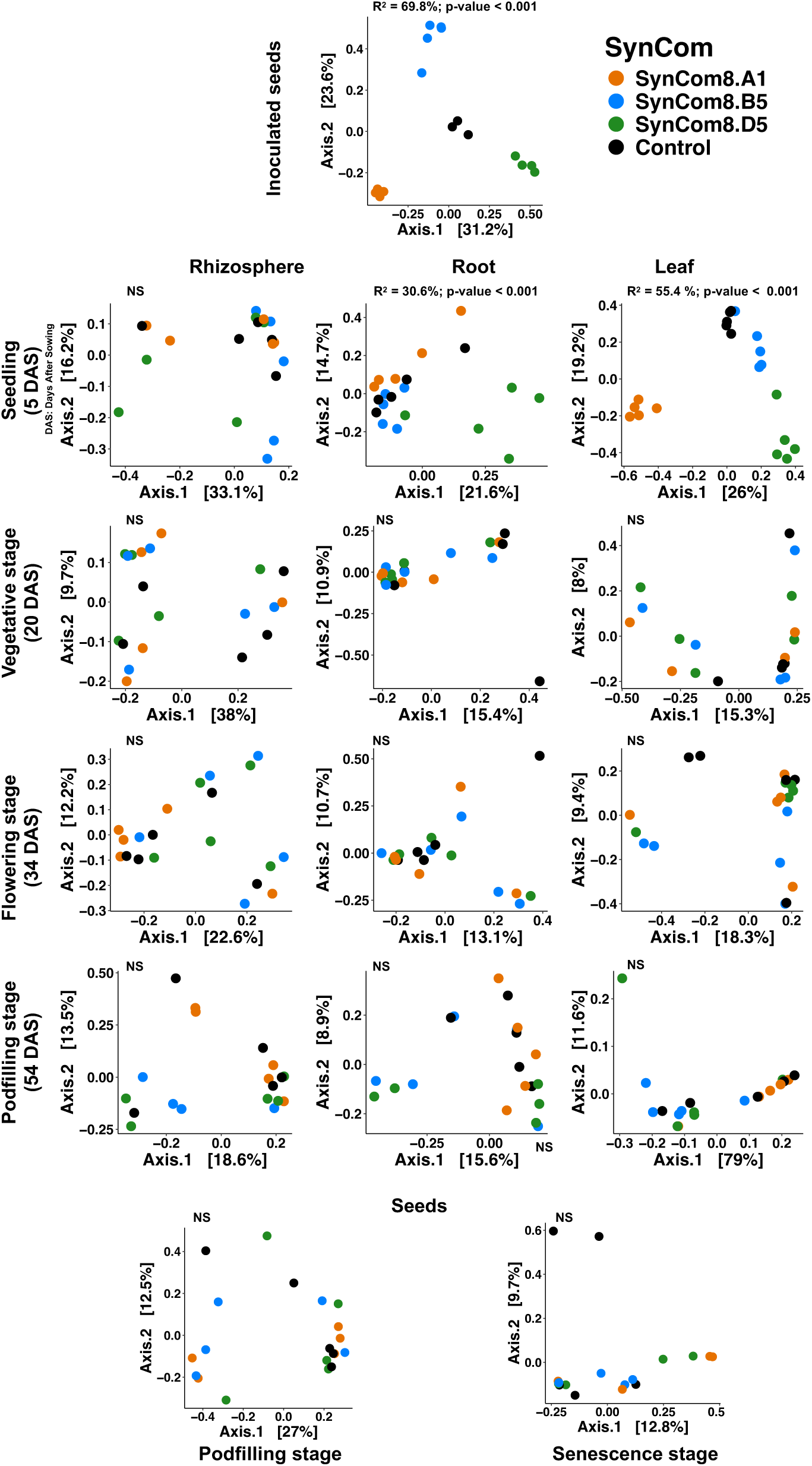
Effect of SynCom inoculation on seed on bacterial communities across stages and compartments. Effect of SynCom inoculation on bacterial communities across plant development stages and compartments visualized through PCoA ordinations based on Bray–Curtis distances. PERMANOVA were used to test the effect of SynCom inoculation on the microbiota structure.

Fungal communities were analyzed targeting the ITS1 region. As for bacteria, the plant stage structured the fungal rhizosphere, root and leaf communities (Figure 7-A, PERMANOVA; R^2^ = 22.5%, 20.7%, and 31.2% respectively, p-values < 0.001). Fungal rhizospheres were dominated by *Leotiomycetes* and *Tremellomycetes*, root communities by *Leotiomycetes*, *Tremellomycetes* and *Agaricimycetes*, and leaf communities by *Leotiomycetes* and *Tremellomycetes* at seedling stage and *Dothideomycetes*, *Tremellomycetes and Eurotiomycetes* for the older stages (Figure 7-B). Seed SynComs inoculation influence only seedling fungal communities, specifically in root and leaf (Figure 8-A, PERMANOVA; Root: R^2^ = 23%, p-value = 0.013; Leaf: R^2^ = 21%, p-value = 0.027). However, even if SynCom did not impact fungal microbiota at a large scale for the other stages (FigS4), differential abundance testing revealed that many fungal genera per stage and compartment were significantly enriched/depleted based on the initial SynCom inoculation (Figure 9-B, p-values < 0.05). For instance, SynCom8.D5 inoculation induced enrichments in the genus *Scheffersomyces* (a Saccharomycetes) and *Meliniomyces*, among others (Figure 7-B, Figure 8-B). The enrichment of *Scheffersomyces* genus was maintained until vegetative stage. Also, *Pseudeurotium* genus was enriched in seedling leaves coming from SynCom8.A1 while it was depleted for SynCom8.B5 (Figure 9-B). Thus, the seed microbiota led to changes in fungal microbiota assembly trajectories over time.

**Figure 7:**
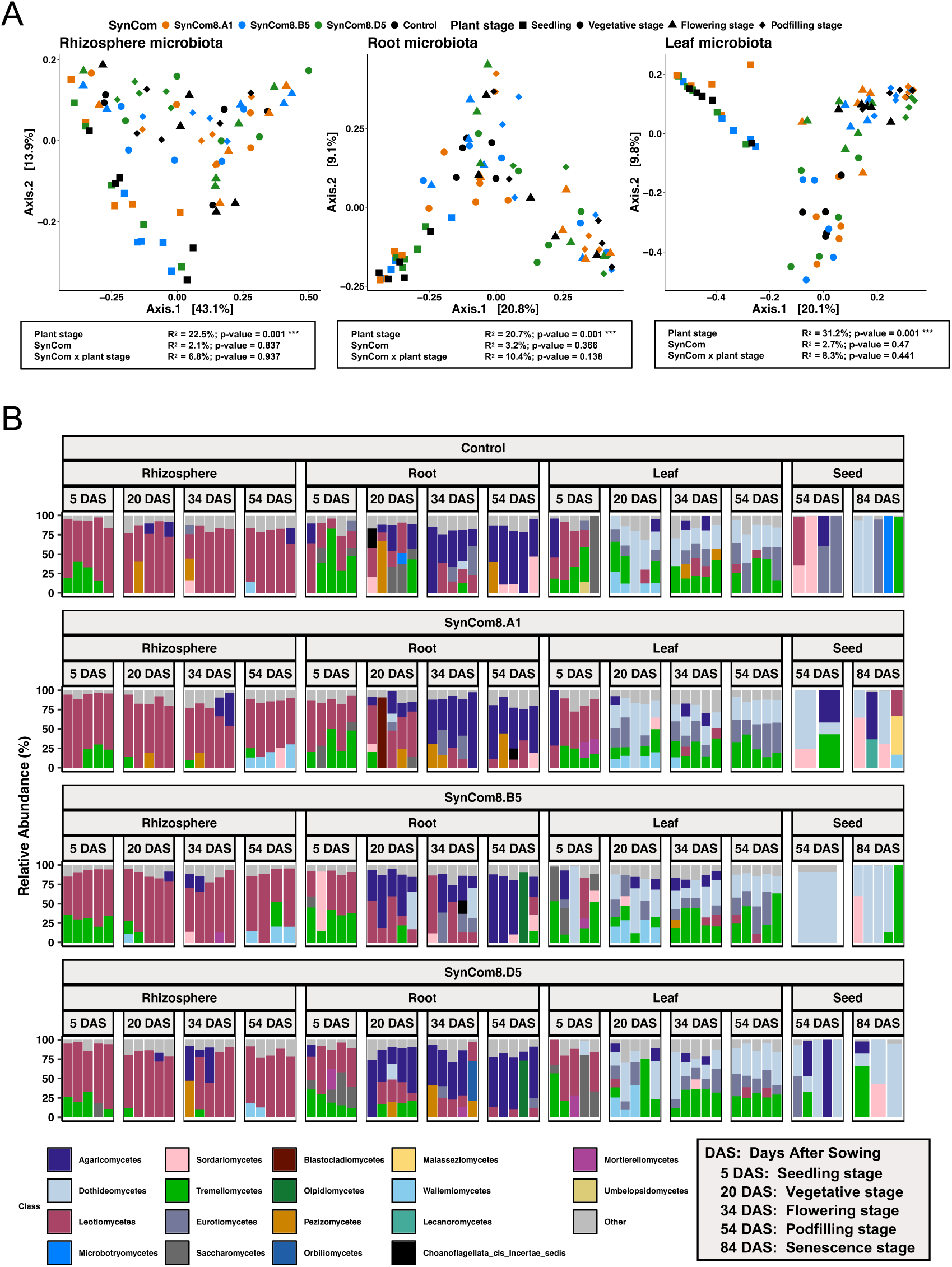
Overview on fungal communities across plant stages and compartments. (A) Effect of SynCom and plant development stages on rhizosphere, root and leaf fungal microbiota structure visualized through PCoA ordinations based on Bray–Curtis distances. PERMANOVA were used to test the effect of plant development stage, SynCom and their interaction for each plant compartment. Samples are shaped based on their plant stage and colored by their initial SynCom composition. (B) Taxonomic profiles of fungal microbiota at order level across stages and SynComs. Taxa representing less than 10% of relative abundance were merged in the “Other” category.

**Figure 8:**
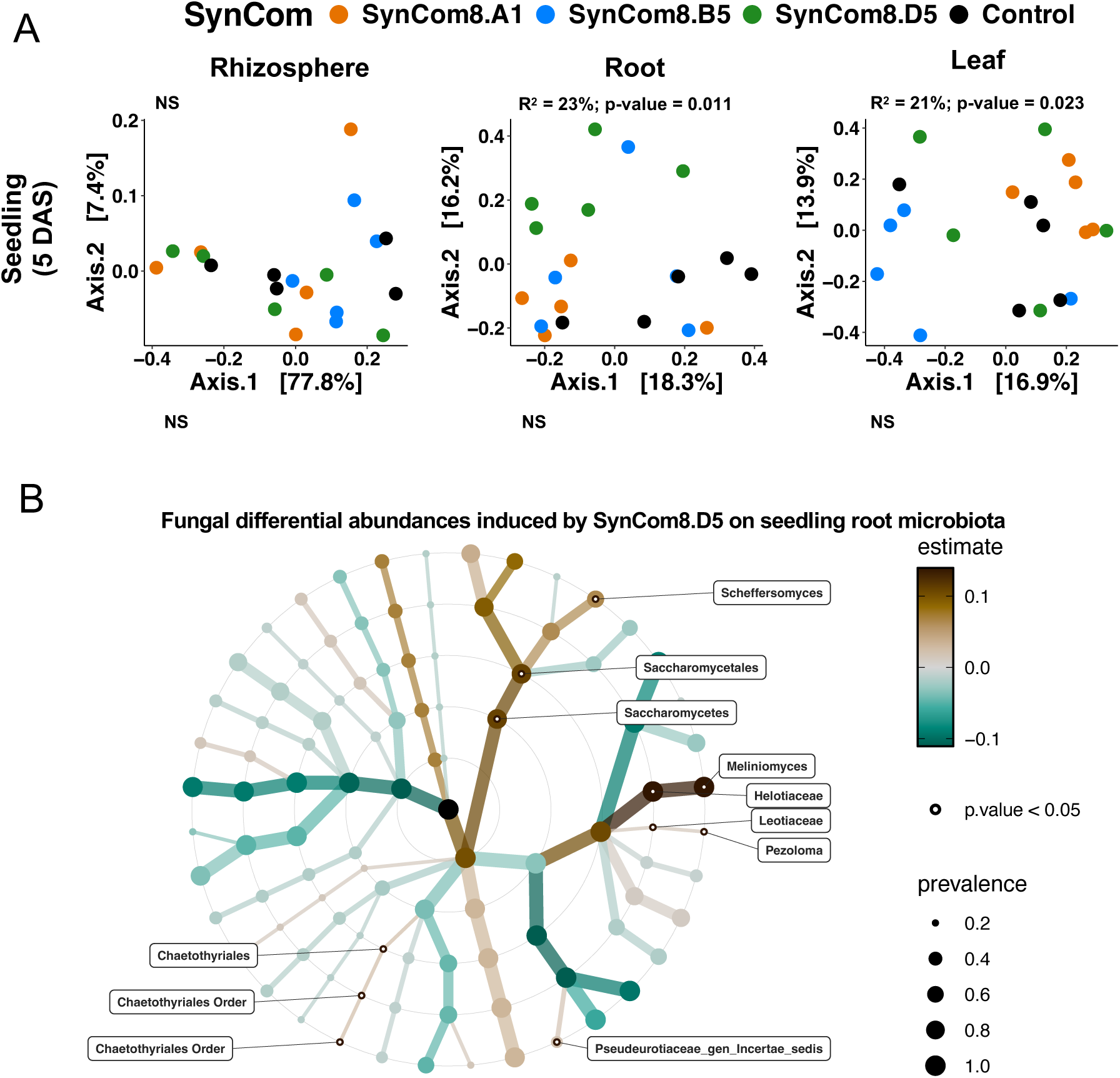
Effect of SynCom inoculation on seed on fungal communities across stages and compartments. (A) Effect of SynCom inoculation on seedling fungal communities visualized through PCoA ordinations based on Bray–Curtis distances. PERMANOVA were used to test the effect of SynCom inoculation on the microbiota structure. Pairwise permanova were made for seedling root and leaf microbiota. The effect of SynCom inoculation on fungal communities for the other stages are displayed in FigS4 as they were showing no significance. (B) Focus on the effect of SynCom8.D5 on seedling root microbiota. Changes in the relative abundance of fungal taxa of root induced by SynCom8.D5 at the different taxa levels using a linear model. Labels of the corresponding taxa were plotted only if p-values < 0.05.

**Figure 9:**
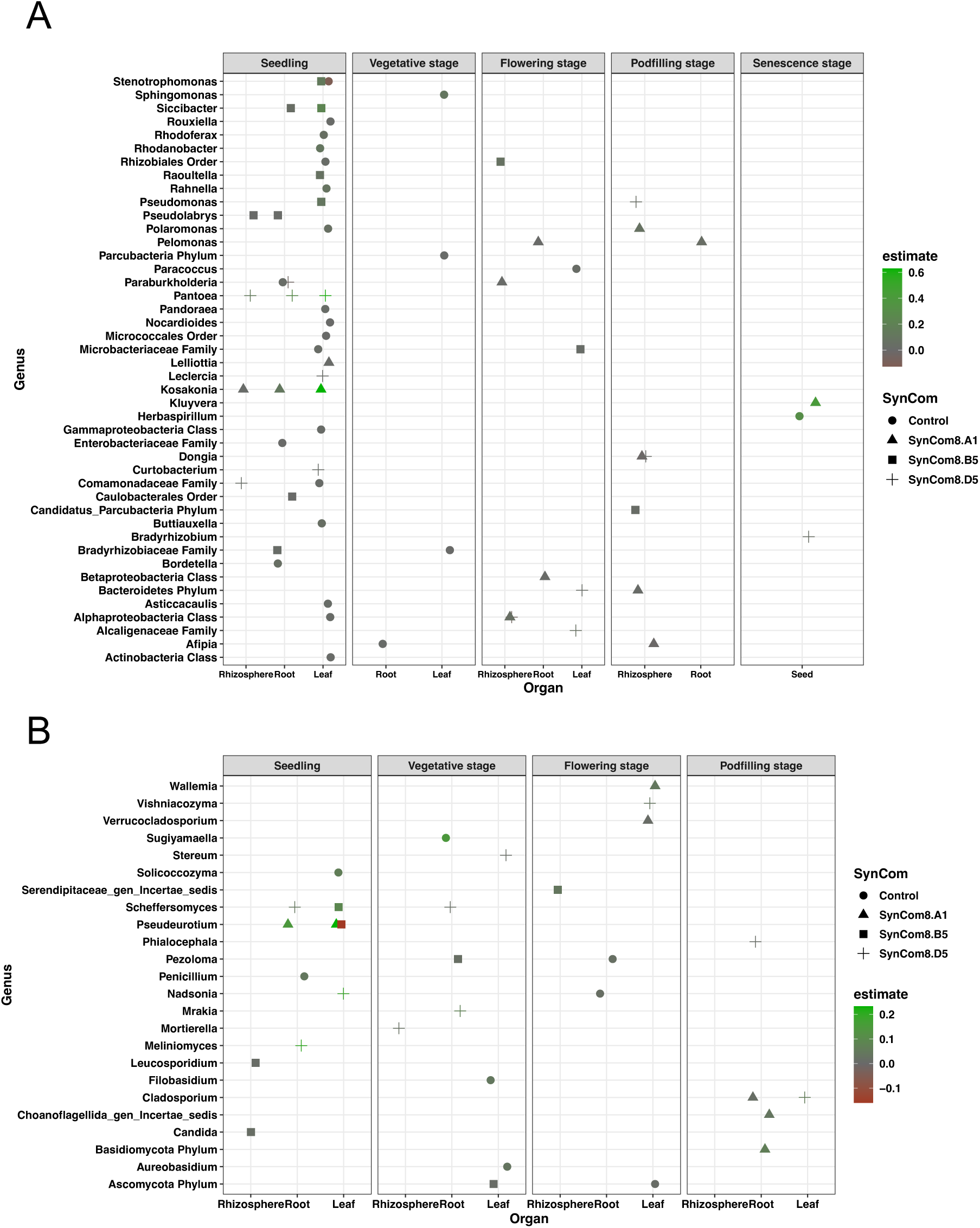
Effect of SynCom inoculations on bacterial and fungal genera across plant developmental stages and compartments. Changes in the relative abundance of bacterial (A) and fungal (B) genus using a linear model. Genus were plotted only when they showed significant (p-values <0.05) increase or decrease above absolute estimate of 0.01. The analysis was not conducted on mature seed for fungi because too many samples presented too few reads.

## Discussion

### Seed microbiota influence on plant microbiota assembly

As other plants, the common bean microbiota composition varies across plant development stages and is structured by compartments (Edwards et al., 2018). Using SynComs we showed that the common bean seed microbiota can be transmitted and contribute actively to the assembly of the seedling microbiota. These results are consistent with other studies showing a high proportion of seed-derived microorganisms in seedling microbiota, whether under conditions of community inoculation (Arnault et al., 2024, Simonin et al., 2023) or natural communities (Abdelfattah et al., 2021, Johnston-Monje et al., 2021). After the seedling stage, the persistence of SynCom members is very low. Only a few strains were detected, such as *Kosakonia sp* (CFBP8986) in the roots at the flowering stage, and *Pantoea agglomerans* (CFBP9000) in various leaf samples up to the flowering stage. The disappearance of SynCom members during plant development is likely related to competition with environmental microorganisms, whose successive waves of dispersal exert propagule pressure on the plant’s resident microbial community (Joubert et al., 2024). The low heritability of the seed microbiota across generations was also described for radish seed and rice (Rezki et al., 2018; Guo et al., 2021). However, other studies showed high seed-to-seed transmission in rice and bean (Kim et al., 2022; Zhang et al., 2022; Sulesky-Grieb et al., 2024). Thus, there is therefore no real consensus on the significance of vertical seed-to-seed transmission. Some studies that have demonstrated high levels of vertical transmission were conducted under consistent conditions across generations, suggesting that reacquisition from the environment is also plausible (Kim et al., 2022).

While the SynComs colonization did not persist after the seedling stage, they induced changes in the overall plant microbiota assembly. Indeed, initial bacterial SynComs induced changes in the fungal composition, highlighting potential interactions between bacteria and fungi (Hassani et al., 2018; Frey-Klett et al., 2011). For instance, SynCom8.D5 induced an enrichment in *Meliniomyces*, a genus already isolated from *Populus* root and is described as a mycorrhizae (Vrålstad et al., 2002; Hambleton et Sigler 2005) or as a root endophyte (Bonito et al., 2016). SynCom8.D5 also increased the relative abundance of *Scheffersomyces* genus in the root at seedling and vegetative stages. Thus, despite disappearing, the seed microbiota could influence the recruitment of secondary microorganisms including symbionts, as previously demonstrated (Ridout et al., 2019; Arnault et al., 2024). Also, seed microbiota of the next generation presented different compositions in some bacterial genus according to initial seed microbiota composition of the first generation. Specifically, uninoculated seeds showed higher proportions of *Herbaspirillum*, a genus that dominated the leaf microbiota. Also, the seeds coming from SynCom8.A1 presented a higher proportion of *Enterobacteriaceae*, specifically in the genus *Kluyvera*. Interestingly, this SynCom8.A1 includes four different *Enterobacteriaceae* strains of different genus (*Lelliottia sp* CFBP8978, *Pantoea agglomerans* CFBP8989, *Kosakonia sp* CFBP8986 and *Siccibacter turicensis* CFBP8990). The relationship between initial SynCom composition and its influence on the seed microbiota of subsequent generations requires further investigation.

The three different SynComs had a greater colonization success on the leaf, highlighting potential niche specificity. Indeed, their relative abundance was greater in leaf, than root and rhizosphere. This finding is also consistent with results obtained in other plant species such as oak seedlings (Abdelfattah et al., 2021), maize (Johnston-Monje et al., 2021), and the shrub *Haloxylon salicornicum* (Laurent-Webb et al., 2023). This niche specialization has already been discussed by Abdelfattah et al. (2022). Also, many studies reported that seed borne taxa are weak root and rhizosphere colonizers (Rochefort et al., 2021; Yang et al., 2017; Leff JW et al., 2017; Green SJ et al., 2006; Ofek et al., 2011; Normander B et al., 2000), as these compartments are mostly colonized by soil microbiota members. Additionally, higher proportions of seed-derived members are found in the rhizosphere when plants are grown in sterile substrate compared to natural soil (Johnston Monje et al., 2016). Thus, niche specialization might drive this partitioning. However, In our case, the relative abundance of a strain on a leaf was correlated with its relative abundance on the root. This correlation suggests that the higher relative abundance of seed-derived strains on leaf is mainly due to priority effect and weaker competition pressure in above ground compartment compared to underground compartment. In other words, because microbial biomass and diversity is higher in underground compartments compared to aboveground compartments, it reduces the chance of SynCom to colonize these underground niches. This hypothesis can be developed in the coalescence framework.

The seedling microbiota assembly can be seen as the coalescence between the seed microbiota and the soil microbiota (Joubert et al., 2024). A coalescence is defined as the mixing between two or more communities in a given environment, with a resulting community displaying different properties and functions from its source communities (Rillig et al., 2015; Joubert et al., 2024). In a coalescence, the respective biomass and richness of each community are a key determinant of the possible outcomes (Rillig et al., 2015, Joubert et al., 2024). Indeed, the more biomass and richness a community has, the more chance it has to “win” the coalescence and dominate the resulting community. For the seed and soil microbiota coalescence, the soil richness and biomass are key factors determining the relative contribution of soil *versus* seed microbiota to the seedling microbiota (Walsh et al., 2021; Arnault et al., 2024). In a previous study, we showed that the more the seed is colonized, the more it colonizes the seedling, which highlights the importance of the mixing ratio to win the coalescence between seed and soil microbiota (Arnault et al., 2024). In the same vein, Arias et al. (2020) showed that transmission of *Xanthomonas vasicola* pv. *vasculorum* is very low under natural conditions (∼2%), whereas it is systematically transmitted to the seedling stage when inoculated at a concentration higher than 10^6^ CFU/mL (Arias et al., 2020). Thus, we could legitimately associate the better colonization of aerial parts by SynCon strains to weaker competition (lower richness observed on leaf), compared with the strong competition that takes place in soil due to the higher biomass and diversity of the soil microbiota.

In conclusion, the seed microbiota colonizes the early stages of plants. In particular, because it benefited from priority effects and weaker competition with microorganisms from the environment, it contributes more to the aerial microbiota of the seedlings. Seed-derived microorganisms are then gradually replaced by continuous colonization of microorganisms from the environment. Still, its early influence can change the trajectory of assembly, favoring specific ecological successions, generating legacy effects even on the new seeds produced.

### Consequences for plant microbiota engineering

For the three SynComs tested, strains showed contrasted transmission capacities and the SynCom composition influenced these colonization capacities. These findings are crucial for future plant microbiota engineering design as not all strains are able to effectively colonize the plant and even high colonizers can be less effective depending on seed and soil community compositions. Despite its transient colonization, the seed microbiota is nonetheless important, as the seed microbiota enhances various aspects of seed physiology (Nelson 2018; Verma et al., 2019; Pal et al., 2022; Kaur et al., 2022; Hone et al., 2021). Thus, the seed-to-seedling transition is of great interest to improve early plant development, limit pathogen colonization and change the trajectory of the plant microbiota assembly through priority effect. However, the low level of strain colonization beyond seedling stage will undoubtedly make it difficult to use seeds as vectors for late-acting biocontrol or biostimulant (Rocha et al., 2019). Finally, from a risk assessment point of view, the low level of colonization of the immediate environment (here the rhizosphere) and the absence of seed-to-seed transmission are advantages for the SynComs tested to limit the dispersal of invasive microbial species, but will have to be studied on a case-by-case basis for each future SynCom developed.

## Conclusion

The goal of this study was to assess the influence of seed microbiota composition on the plant microbiota assembly across compartments and stages during an entire plant life cycle. Bacterial SynComs were reconstructed using seed isolated strains to manipulate the seed microbiota and control its natural variability. By doing so, we set up different seed microbiota compositions as primary inoculum for the plant microbiota assembly. We showed that the seed microbiota has a transient colonization success to seedlings before being replaced by environmental microorganisms. However, this transient colonization is nonetheless important, as it can influence the recruitment of bacterial or fungal microorganisms that might shape the plant microbiota assembly through priority effect. These findings highlight the importance of the seed microbiota as a primary inoculum of the plant. We hope that further efforts will be made to better characterize this important compartment, which constitutes both a starting point and an end point of the plant microbiota.

## Supplementary figures

**Figure S1:**
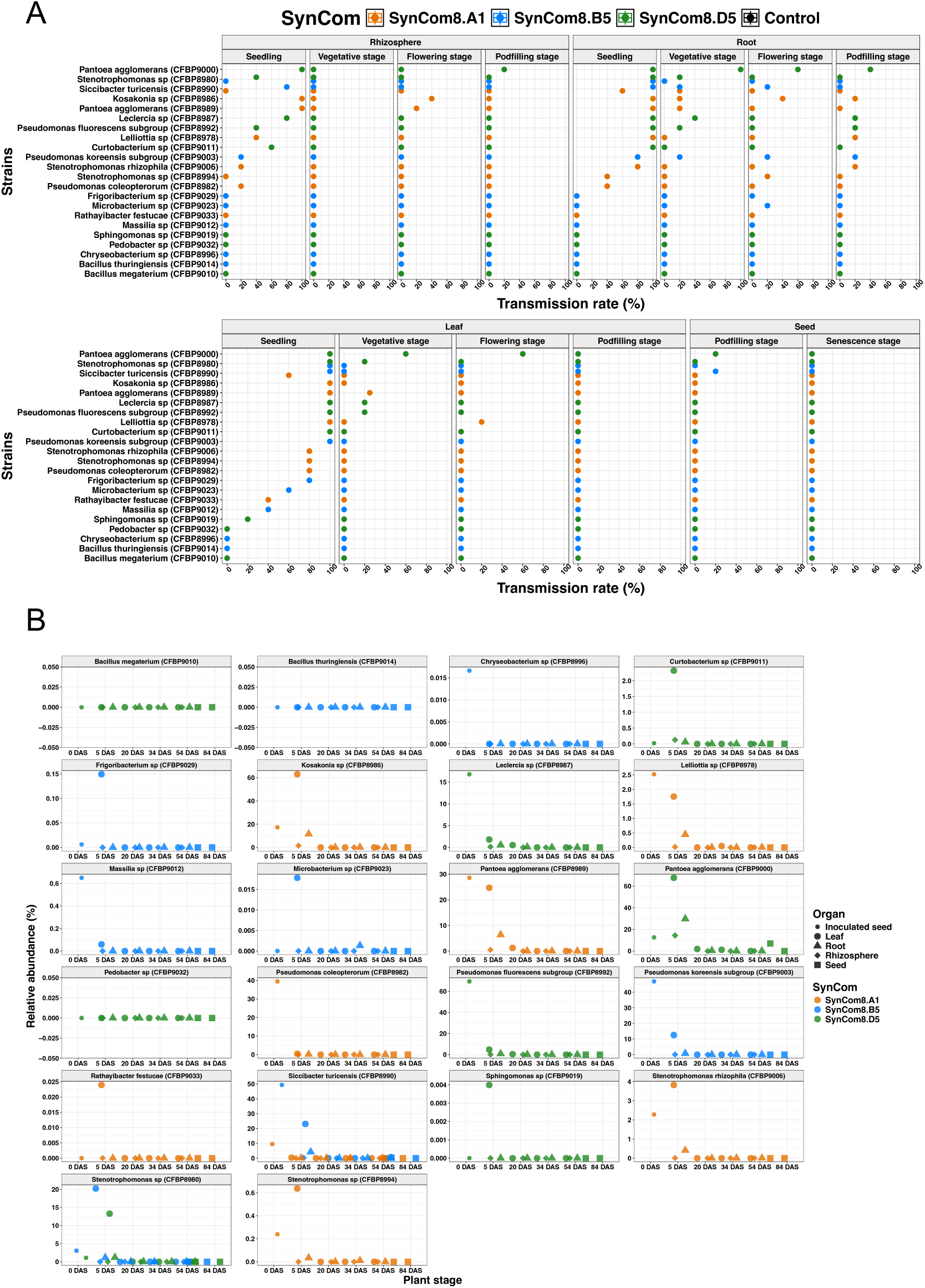
Inoculated strain colonization success over development stages and compartments.

**Figure S2:**
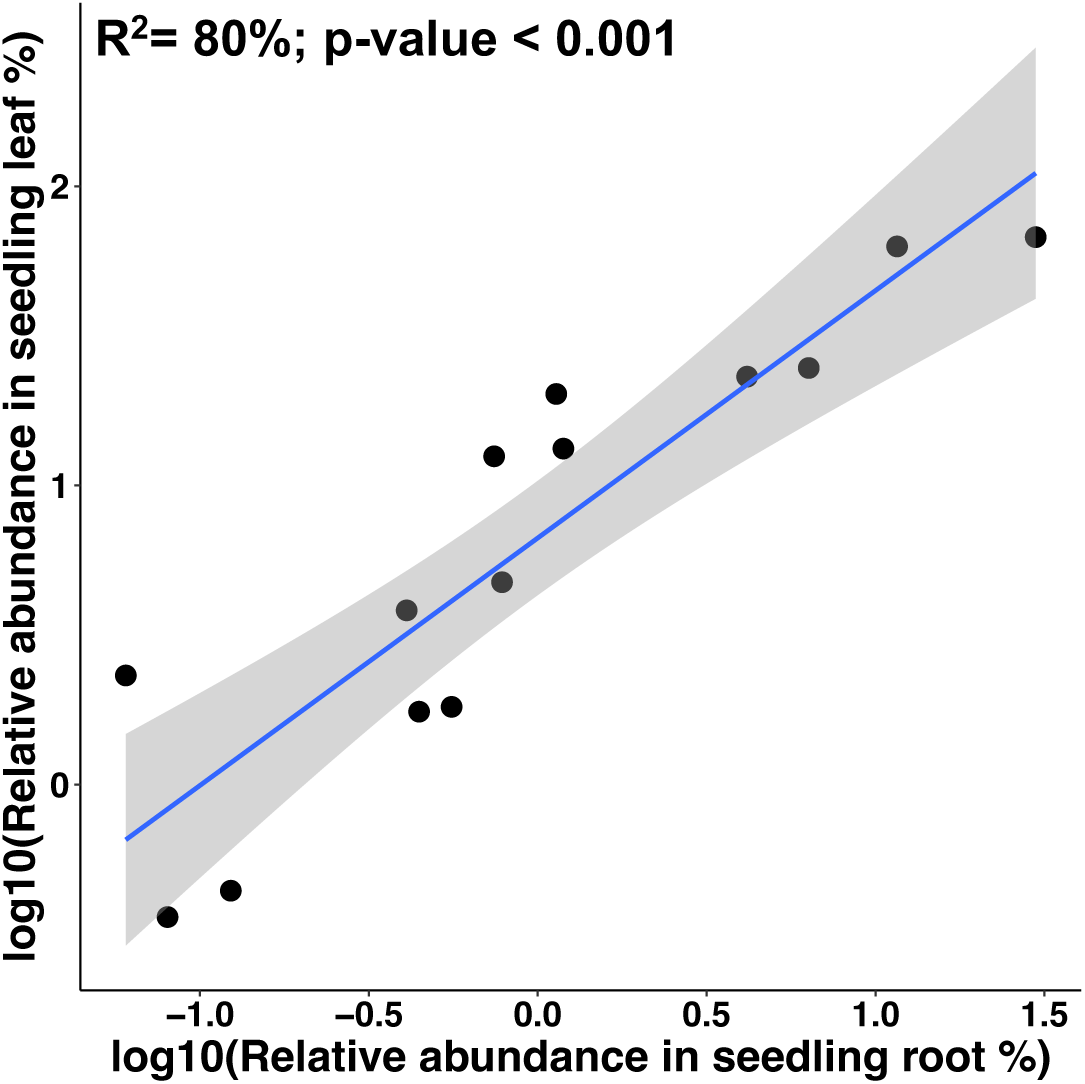
Linear regression between the relative abundance of a strain in the leaf and its relative abundance in the root.

**Figure S3:**
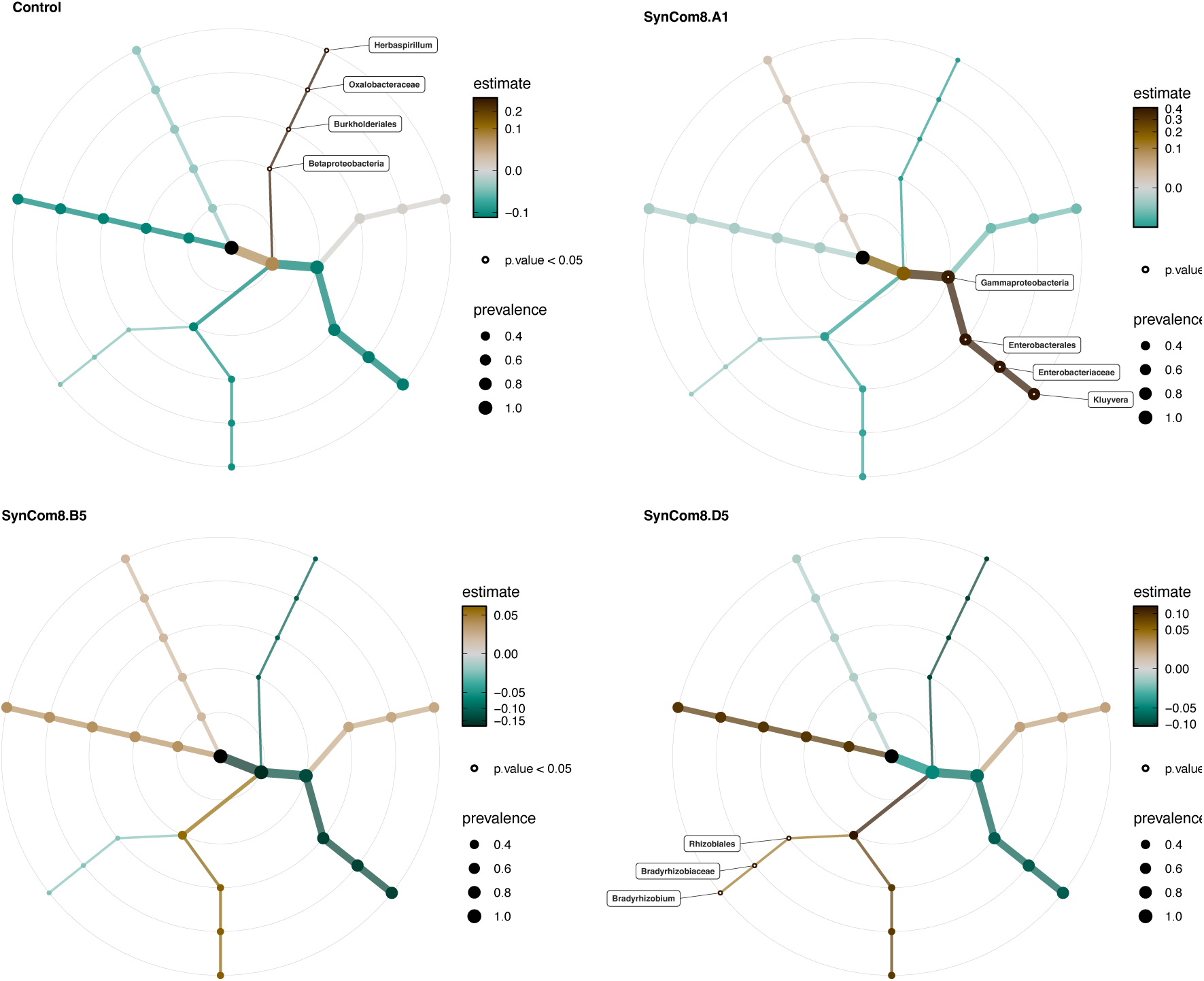
Differential abundance testing of the bacterial community on mature seed of the next generation.

**Figure S4:**
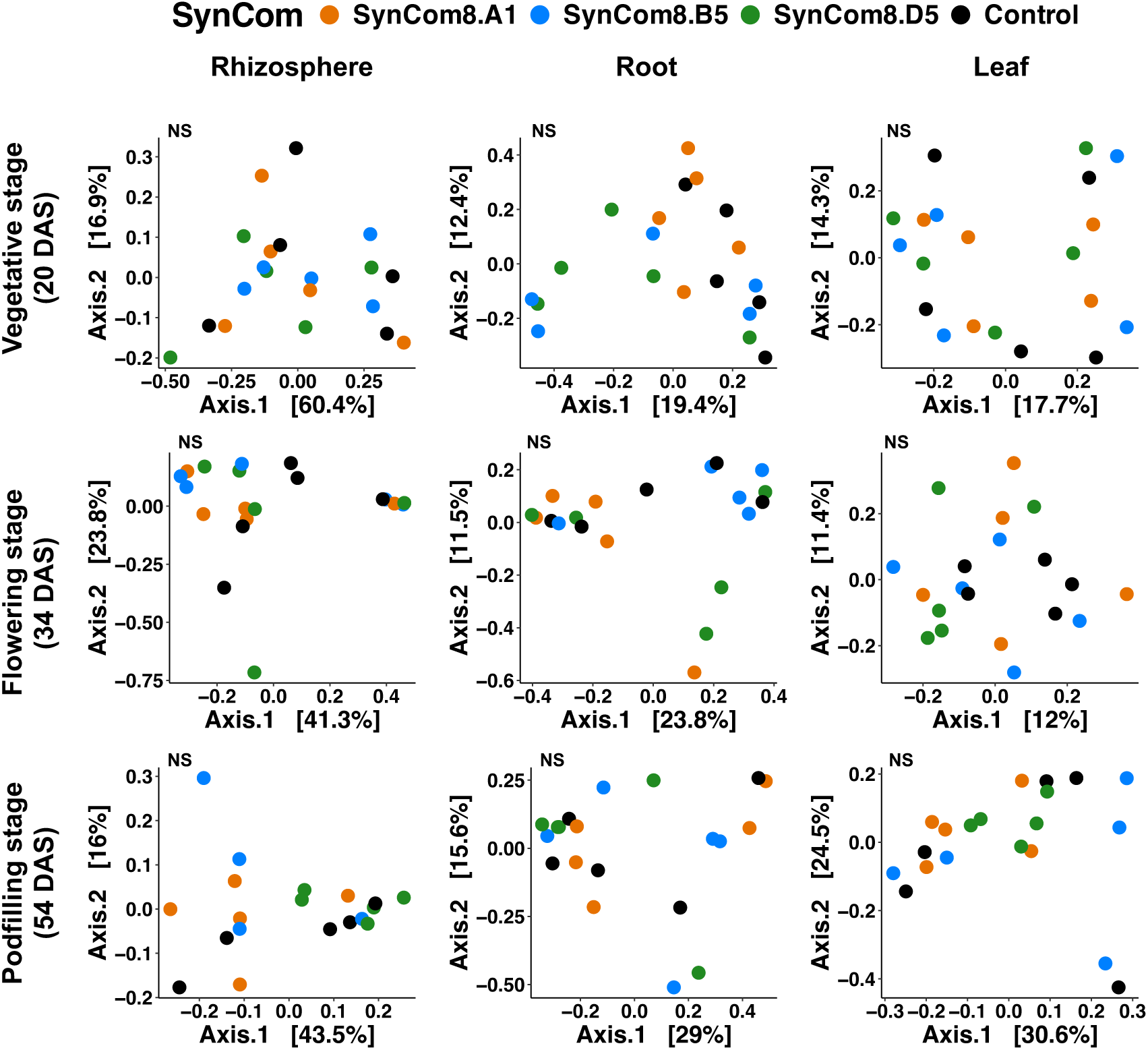
Effect of SynCom inoculation on seed on fungal communities across stages and compartments. Effect of SynCom inoculation on fungal communities across plant development stages and compartments visualized through PCoA ordinations based on Bray–Curtis distances. PERMANOVA were used to test the effect of SynCom inoculation on the microbiota structure. Pairwise permanova were made for seedling root and leaf microbiota.

